# A Systematic Review and Meta-analysis on the Transcriptomic Signatures in Alcohol Use Disorder

**DOI:** 10.1101/2022.12.19.521027

**Authors:** Marion M Friske, Eva C Torrico, Maximilian JW Haas, Anna M Borruto, Francesco Giannone, Andreas-Christian Hade, Yun Yu, Lina Gao, Greg T Sutherland, Robert Hitzemann, Mari-Anne Philips, Suzanne S Fei, R Dayne Mayfield, Wolfgang H Sommer, Rainer Spanagel

## Abstract

Alcohol use disorder (AUD) is a complex mental health condition. Currently available clinical treatments exhibit limited efficacy and new druggable targets are required. One promising approach to discover new molecular treatment targets involves the transcriptomic profiling of brain regions within the addiction neurocircuitry, utilizing animal models and post-mortem brain tissue from deceased AUD patients. Unfortunately, such studies suffer from large heterogeneity and small sample sizes. To address these limitations, we conducted a cross-species meta-analysis on transcriptome-wide data obtained from brain tissue of AUD patients and animal models. We integrated 36 cross-species transcriptome-wide RNA-expression datasets with an alcohol-dependent phenotype vs. controls, following the PRISMA guidelines. In total, we meta-analyzed 1,000 samples – 502 samples for the prefrontal cortex (PFC), 318 nucleus accumbens (NAc) samples, and 180 amygdala (AMY) samples. The PFC had the highest number of differentially expressed genes (DEGs) across rodents, monkeys, and humans. Commonly dysregulated DEGs pointed towards enrichment in inflammatory responses and alterations in BBB-regulatory mechanisms in astrocytes, microglia and endothelial cells. Gene set enrichment analysis further showed that MAPK/ERK-signaling plays a critical role in AUD and especially in monkeys *Dusp4* as a major inhibitor of the MAPK pathway may be a main driver of these pathway alterations. Our data also suggest that the transcriptomic profile in the NAc is less vulnerable to the maintenance of AUD. Finally, we provide a combination of DEGs that are commonly regulated across different brain tissues as potential biomarker for AUD. In summary, we provide a compendium of genes, signaling pathways, and physiological and cellular processes that are altered in AUD and that require future studies for functional validation.

## Introduction

Alcohol is one of the most widely consumed psychoactive drugs, globally round 2.3 billion adults drink alcohol at least annually. ^1^ However, alcohol consumption is also associated with many harms, especially when used chronically Alcohol Use Disorder (AUD) may develop, a complex multifaceted human disease that constitutes a major public health problem, representing a complex, multifaceted human disease that poses a significant public health challenge, an extensive economic burden, and imposes a considerable human toll ^2^. AUD is associated with numerous comorbidities, including major depressive disorder, antisocial and borderline personality disorders and other psychiatric conditions, liver and many other organ injuries, as well as numerous cancer types ^3^. Even though alcohol is acting systemically on a variety of organs, the origin of AUD is primarily the brain, which is why AUD is considered as a brain disease ^4, 5^.

A key goal of AUD research is to unravel the underlying molecular changes that contribute to disease progression, maintenance and relapse after abstinence and to pinpoint essential molecular targets for the development of new medications ^6–8^. Numerous preclinical studies have shown that chronic alcohol consumption disrupts multiple molecular pathways in various brain regions ^9–11^. As a result of these molecular alterations, alcohol affects the activity of neuronal circuits ^12^, especially in the extended reward system and addiction neurocircuitry ^13, 14^. Together, these mechanisms produce long-lasting cellular and network adaptations in the brain that in turn can drive the development and maintenance of AUD and relapse behavior.

Although preclinical microarray and RNA-sequencing studies in different model organism such as mice, rats and monkeys provide valuable brain transcriptomic information upon chronic alcohol consumption, our understanding of how effectively these findings translate to the human AUD condition remains limited. A major limitation of transcriptome-wide approaches in animal models is the often inadequate sample size, leading to insufficient statistical power and unreliable findings ^15^. Additionally, inter-study variability due to e.g., breeding facility, laboratory conditions, different strains and experimental protocols, leads to diverse outcomes among published studies. These limitations hinder the translatability of preclinical research and underscore the need for optimizations to yield meaningful conclusions ^16^.

Researchers have also attempted to explore the transcriptomic profiles of post-mortem brain tissue from deceased AUD patients ^17–20^. However, these post-mortem brain studies suffer from large variation in the quality of the provided brain tissue and the large heterogeneity of AUD as such. The heterogeneity results from different genetic backgrounds and personal life histories such as early life adversity, age, sex, infectious history, tobacco and poly-drug use, and many other factors. Hence, very large cohorts are required to achieve sufficient statistical power and to allow the identification of generalizable and reproducible transcriptome-wide signatures.

Here, we merged multiple publicly available and also some unpublished datasets in a cross-species approach with the idea of overcoming some of the aforementioned limitations and to identify alterations in gene expression profiles and regulatory pathways that could previously not been identified due to low statistical power and heterogeneity of the experimental and pathological condition. For doing so, we meta-analyzed data from different microarray and RNAseq-platforms retrieved from different brain areas of the addiction neurocircuitry of AUD patients, from alcohol-dependent rats, mice and monkeys as well as from appropriate controls.

Although no animal model of addiction fully mimics the complex human condition, certain animal models are better suited to investigate molecular alterations that might be relevant for the human condition. Here, we focused our attention on the post-dependent animal model where rats and mice undergo chronic intermittent alcohol exposure. This animal model mimics the natural development of AUD in humans characterized by several intermittent drinking episodes interspersed with withdrawal periods ^21^. Importantly, chronic intermittent alcohol exposure results in robust and stable behavioral symptoms of alcohol dependence that persist during prolonged abstinence and is characterized by long-lasting molecular neuroadaptations that mediate such symptoms ^22^. Based on the number of studies identified through our initial systematic literature screening, the meta-analyses of this study are concentrated on three critical important areas of the addiction neurocircuitry, namely the prefrontal cortex (PFC), nucleus accumbens (NAc) and amygdala (AMY). Other brain sites could not be considered for meta-analyses due to low study and sample numbers.

The here presented convergent cross-species findings on differentially expressed genes (DEGs) is providing a foundation for further functional research into understanding the mechanisms of AUD in the brain and for developing subsequently new therapeutic targets.

## Materials and Methods

### Systematic Literature Screening

The systematic literature screening was performed according to the PRISMA Guidelines (Page et al., 2021). The protocol for this review has been registered at PROSPERO and Open Science Framework (Open Science Framework: https://doi.org/10.17605/OSF.IO/TF8R4; PROSPERO: CRD42020192453). Briefly, two researchers independently screened the literature on PubMed and EMBASE with the predefined keywords that were determined to specify four main categories of interest: alcohol/ethanol AND species-specific & model-defining terms AND RNA-specific terms AND brain (**Fig 1A**, for detailed keyword information see Supplementary).

**Fig. 1:**
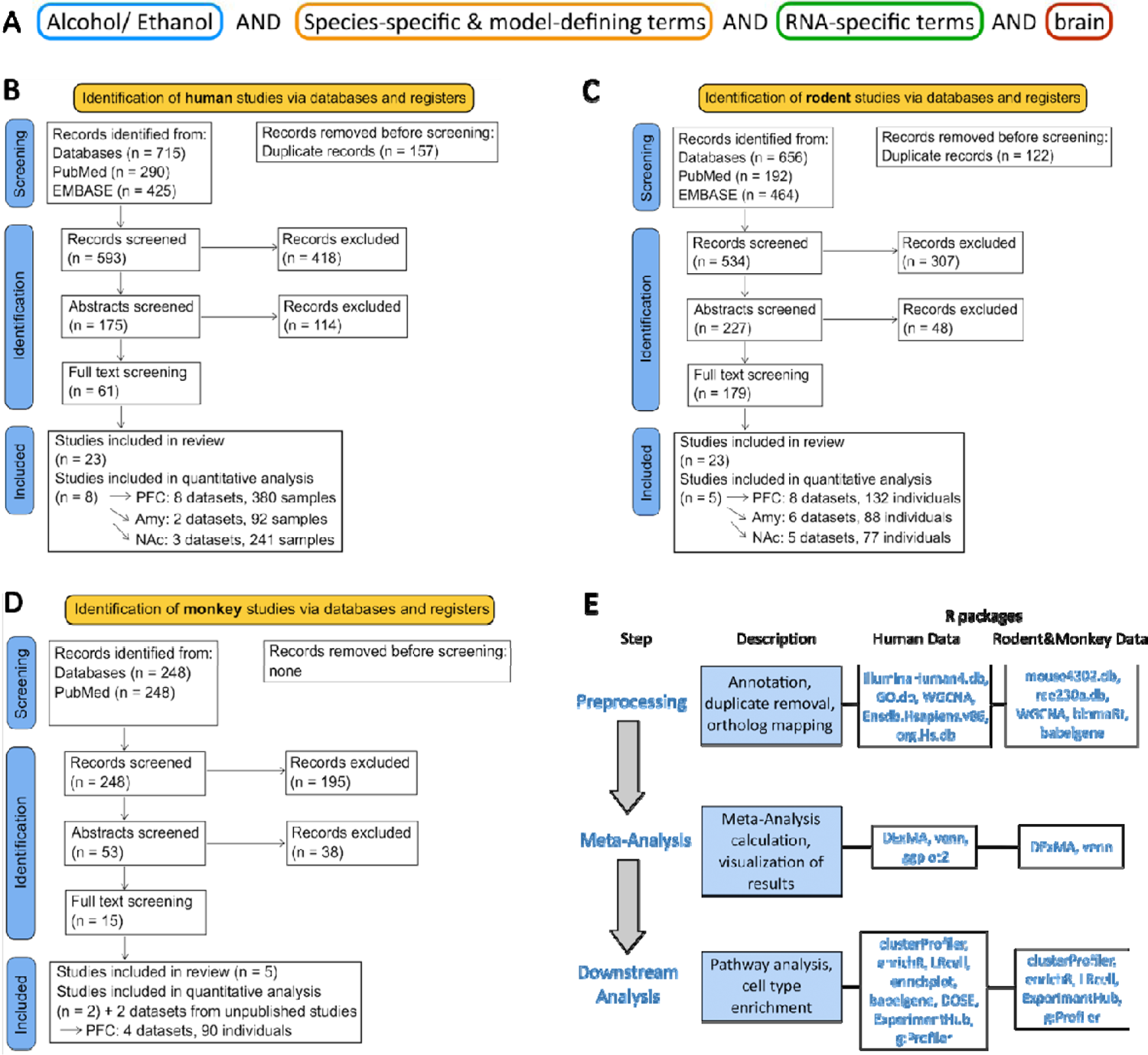
Systematic literature screening of PubMed and EMBASE to retrieve transcriptome-wide expression datasets derived from (B) post-mortem brain tissue of deceased AUD patients and controls, (C) from post-dependent rodents, and (D) from monkeys that had long-term intermittent voluntary alcohol consumption. (A) General structure for the keyword design for systematic literature research (B) PRISMA workflow for the screening of human studies. Before the screening procedure, duplicates due to overrepresentation in the two databases screened were removed by using EndNote. The template for overview of the screening procedure and the resulting studies was taken from www.prisma-statement.org. Eventually, we identified ten human postmortem studies matching our criteria ^17–20, 23–28^ (C) PRISMA workflow for the screening of rodent studies resulted in five studies to include ^29–33^ (D) PRISMA workflow for the screening of monkey studies identified two studies to include in the meta-analysis ^34, 35^. In addition, unpublished data were kindly provided by Dr. Kathleen A. Grant and Dr. Suzanne Fei. (E) General workflow of the meta-analysis pipeline with the respective packages used for the statistical software R.

The literature screening was conducted using the SysRev screening tool, which was developed by the CAMARADES research group at the University of Edinburgh (www.syrf.org.uk). Following each screening step (title/abstract and full text), the researchers compared the respective resulting studies. In case of disagreement, a third, independent expert in the field chose whether the study was included or excluded for the next screening step. The general structure of the predefined keywords, as well as the complete PRISMA flow diagrams with the resulting number of studies for each screening step for the rodent, monkey and human study screening is depicted in **Fig.1**. The screening and data extraction yielded a substantial number of studies for the PFC, NAc and AMY, enabling us to conduct a comprehensive meta-analysis.

### Inclusion and exclusion criteria for human, rodent, and monkey studies

Only studies providing transcriptome-wide RNA-expression datasets, such as microarray or RNA-Seq experiments, were considered for the meta-analysis. The following inclusion and exclusion criteria were applied during screening. Inclusion criteria for rodent studies were: rodents (mice/rats), alcohol exposure (≥ 2 weeks), post-dependence (≥ 3 days abstinence prior to death), transcriptome-wide RNA-expression data (microarray/RNA-Seq) from brain tissue, and controls without alcohol intake. Studies were excluded in case of: adolescence, prenatal alcohol exposure, too short exposure/abstinence times, or additional treatments. For human post-mortem studies, the inclusion criteria were: human post-mortem tissue, diagnosed AUD (DSM-IV), transcriptome-wide RNA-expression data (microarray/RNA-Seq) from brain tissue, non-AUD controls. Studies were excluded in case of other psychiatric comorbidities or brain injury. For screening of monkey literature, the following inclusion criteria were applied: non-human primate, transcriptome-wide RNA-expression data (microarray/RNA-Seq) from brain tissue, alcohol-naive controls. Studies were excluded if monkeys received any additional treatments or were subject to prenatal alcohol exposure. The summary statistics, which included the comprehensive profiles of all detected genes from the sequencing experiments, were either obtained by downloading from the supplementary materials of the original publications or by reaching out to the corresponding authors to request the respective datasets. Details of the included studies are listed in the Supplementary.

As is customary in systematic reviews and meta-analyses, we aimed to assess the risk of bias (RoB) for the original studies. Unfortunately, inadequate reporting of the conducted experiments hindered our ability to access the complete information required for a proper RoB assessment.

### Meta-analysis approach for the three species

The aim of this project was to provide a highly comprehensive input dataset that consists of as many original datasets as possible matching our criteria. Due to data availability constrains, the analysis was performed using summary statistics. Accessing the original data was not feasible, as many of the studies had been conducted some years ago. Despite attempts to contact most of the corresponding authors, the raw data could no longer be retrieved. To maximize the number of studies to combine and as the data was derived from different platforms and some lacked standard error distribution information, a p-value combination approach was chosen. This type of meta-analysis has been previously used for transcriptome-wide approaches ^36–38^, especially when data from different platforms are combined, demonstrating its suitability for our study. Standard p-value combination meta-analyses fail to account for contradictory expression patterns, while Stouffer’s p-value combination approach is uniquely enabling the account for contradictory expression patterns by using one-tailed p-values as input ^39^. The meta-analyses were performed by transforming the two-tailed p-values from the original publications into one-tailed p-values according to the observed effect direction in the respective studies as previously described ^36, 40, 41^. Subsequently, two separate meta-analyses were conducted per species using the R package DExMA (https://www.mdpi.com/2227-7390/10/18/3376), one on left-sided p-values testing for downregulation and one on right-sided p-values testing for upregulation for each gene. Both results were then combined by excluding the higher p-value for each gene and a false discovery rate was calculated. Stouffer’s method additionally provides the possibility to include weights, which was here considered as the square root of the respective sample size for each original study, previously shown to improve power substantially ^42^. While this approach did not provide a pooled estimator for effect size, the median fold-change across studies for each gene was calculated to indicate the direction of the observed effect. For the analysis, we excluded non-protein coding genes.

### Assessment of robustness and heterogeneity

To assess the robustness of meta-analysis results, we conducted leave-one-out-meta-analyses (LOO-MA) on the human PFC datasets. In this analysis, we systematically excluded the study with the highest sample size and the largest number of reported DEGs, which is Kapoor et al. (2019).

As an indicator of robustness, we examined the overlap among the results obtained from the main meta-analysis (list of DEGs with FDR < 0.1), the LOO-MA results, and the reported DEGs from Kapoor et al. (2019). To assess the heterogeneity of DEGs across the studies included in our analysis and their respective meta-analyses, we created Venn diagrams illustrating the intersection of the identified DEGs (FDR < 0.1).

The majority of included rodent studies as revealed by our systematic literature screening, predominantly consisted of mouse studies – depending on the brain region the range was 75% to 83% of the total. Therefore, in order to understand the species-specific impact on the meta-analysis outcomes for post-dependent rodents and to assess heterogeneity across species, we conducted subgroup analyses by categorizing the original studies based on the species involved, specifically mice and rats. Subsequently, we performed separate meta-analyses within the subgroups and compared their results with the DEGs obtained from the combined meta-analysis that considered both rodent species together in Venn diagrams.

### Gene Set Enrichment Analysis

To identify potential pathways and biological functions associated with the DEGs identified in the meta-analyses, we carried out gene set enrichment analysis (GSEA) utilizing GO-terms, Reactome-pathways and KEGG-pathways. This analysis was performed using the R packages ClusterProfiler ^43^ and g:Profiler ^44^. As input for ClusterProfiler, we ranked DEGs based on the formula -log10(p-value) multiplied by the median fold-change. Additionally, we performed cell-type enrichment for the PFC in the human and rodent datasets using the R package LRcell ^45^. To explore protein network associations, we employed STRING to estimate the level of connectivity of the identified DEGs on the protein level.

## Results

### Meta-analysis of the human PFC results in DEGs mainly involved in neuroimmune functions

For the human post-mortem PFC, eight datasets comprising 380 samples were included (**Fig. 1**). The meta-analysis resulted in a total number of 1,945 DEGs with FDR<0.05 (2,803 with FDR>0.1), whereby 78% (n=1,525 DEGs) of the transcripts were up- and 22% (n=421 DEGs) were down-regulated (**Fig. 2A, B**; Supplementary). The top 10 DEGs are mainly involved in neuroimmune functions e.g., *SELE*, *IL1R2*, *CSF3*, and cell cycle regulation e.g., *FOSL1*, *MMP19* and *HSPA6*. STRING analysis was performed to identify the most inter-connected genes among the significant DEGs. For the human PFC, the genes with the highest number of connective nodes were *TP53* (n(nodes)=243), *ACTB* (n(nodes)=242), *MYC* (n(nodes)=193), *JUN* (n(nodes)=155), and *MAPK3* (n(nodes)=150) (Supplementary). Further assessment of the enrichment of the determined DEGs in specific pathways by GSEA resulted in the dysregulation of neuroimmune signaling pathways, cell cycle and cancer-associated pathways, such as interleukins e.g., IL-1, 4, 10, 13, and 17, TNF signaling components, as well as MAPK and ERBB signaling pathway. Cell type enrichment analysis resulted in significant enrichment of the identified DEGs in endothelial cells (FDR<9e-12) and astrocytes (FDR<3e-06) (Supplementary).

**Fig. 2:**
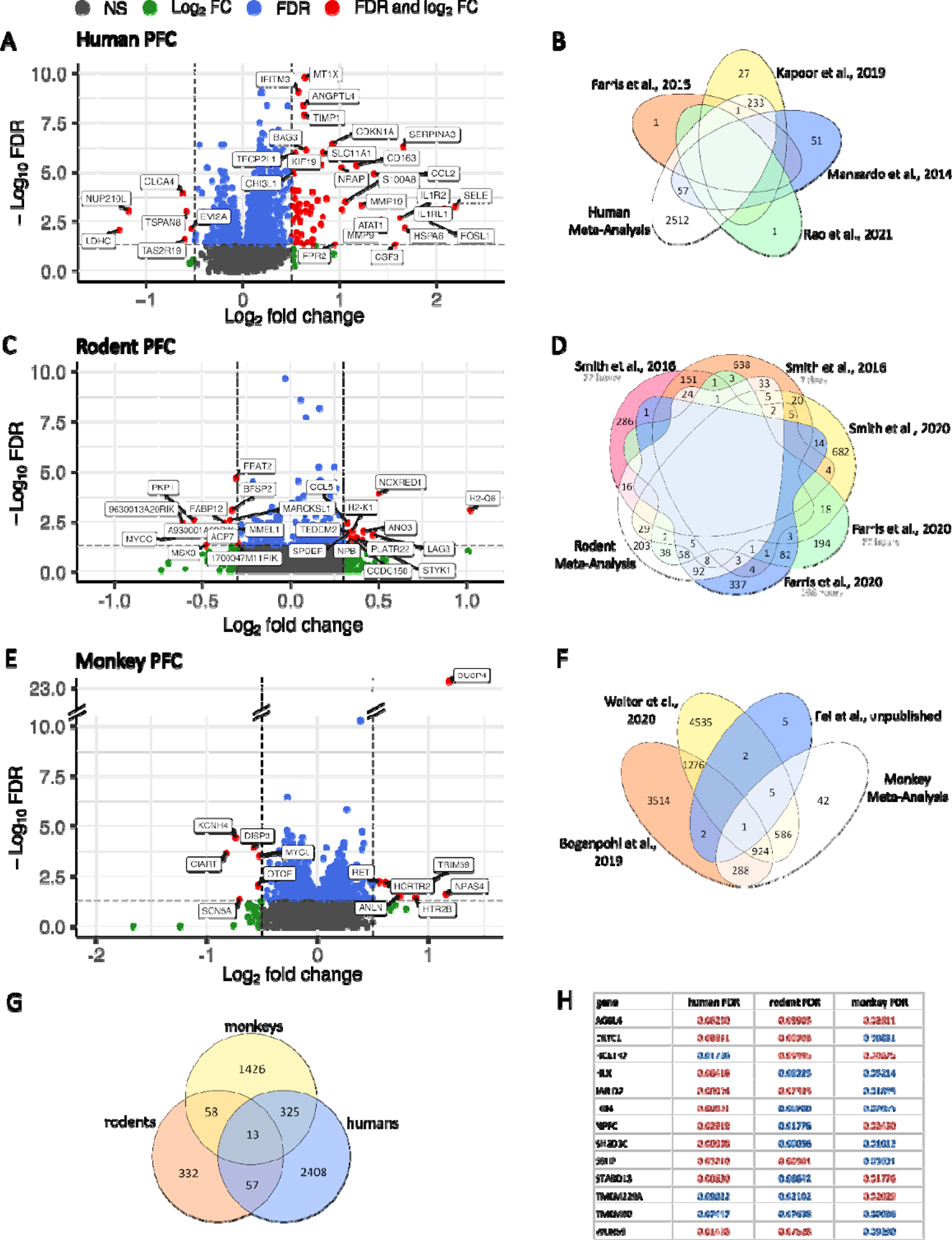
Meta-Analyses of the transcriptome-wide gene expression data derived from the PFC of humans (A, B), rodents (C, D) and monkeys (E, F) identified by species-specific Stouffer’s p-value combination with FDR<0.05 as threshold for significantly altered transcripts (DEGs). Transcripts with FDR<0.05 and log2 fold-change (FC)>0.5 for humans and monkeys and FC>0.3 for rodents are highlighted in red; DEGs with FDR<0.05 and FC<0.25 are highlighted in blue; DEGs with FDR>0.05 and FC>0.25 are highlighted in green; DEGs with FDR>0.05 and FC<0.5 or 0.3 respectively, are highlighted in gray. The Venn diagrams describing the rodent (D) and monkey (F) studies include the time span of last alcohol experience and time point of death as indicated in light gray below the respective study. (G) Venn diagram depicting the cross-species comparison considering DEGs with FDR<0.1 across the human, rodent and monkey meta-analysis results. (H) Thirteen DEGs were detected as significantly altered in the PFC across all three species with two transcripts being dysregulated in the same direction - AGBLA4 and TMEM80. FDR values depicted in blue represent down-regulated transcripts, while FDR values in red stand for up-regulated DEGs.

### Meta-analysis of the rodent PFC results in DEGs that are also enriched in neuroimmune system-related pathways

Systematic literature screening and data extraction of the rodent studies led to 8 datasets including 132 subjects for the meta-analysis of PFC data (**Fig. 1**). The meta-analysis resulted in 299 DEGs with FDR<0.05 (520 DEGs with FDR<0.1) and a distribution of 22% (n=65 DEGs) up-regulated and 78% (n=234 DEGs) down-regulated transcripts (**Fig. 2C, D**; Supplementary). When comparing the common DEGs (FDR<0.05) of the human and the rodent PFC, 14 DEGs are dysregulated in the same direction and are mainly playing a role in neuronal signaling (*Clstn2*, *Rapgef2*, *Tnr*, *Ap2a1*) and neuroimmune pathways (Fkbp5, *Zc3hav1*). When considering a significance threshold of FDR<0.1, which results in 70 DEGs that appear in both, the rodent and the human meta-analysis, around 50% of the transcripts are dysregulated in the same direction. On the level of dysregulated pathways, there are also enriched pathways for both species to note. In both, humans and rodents GSEA points towards an increased impairment in neuroimmune pathways. As described above, in humans the pathways involving interleukin signaling were significantly dysregulated as pointed out by GO-Term, as well as KEGG-pathway and Reactome analysis (Supplementary). In the rodents, T cell regulation pathways including the differentiation, activation and regulation are among the most significant results (Supplementary). Furthermore, the GO-Term GSEA results describe primarily altered pathways of immune regulation e.g., “cytokine production”, “IL-2 production”, “natural killer cell mediated immunity”, and “leukocyte mediated immunity” (Supplementary). When observing the DEGs resulting from the meta-analysis of the rodent datasets, the most significantly altered genes are involved in neuronal signaling e.g., *Ano3*, *Prepl*, *Marcksl1*, and *Btbd8*, immune-regulation e.g., *Lag3*, *Pkp1*, and *H2-q6*, and cancer-related pathways e.g., *Atl1*, *Styk1*, *Spdef*, and *Frat2*. When observing the highest inter-connected transcripts via STRING, *Src* (n(nodes)=39), *Jun* (n(nodes)=26), *Ubxn7* (n(nodes)=25), *Hspa8* (n(nodes)=21), and *Yes1* (n(nodes)=21) had the highest number of nodes.

### Meta-analysis of the monkey PFC results in DEGs that are enriched in cell signaling and MAPK cascade-related processes

For the monkey PFC, systematic literature screening identified two datasets. In addition, we retrieved unpublished data kindly provided by Suzanne Fei, Robert Hitzemann, and Kathleen Grant, which resulted in a final number of four studies comprising 90 individuals (**Fig. 1**). Meta-analysis resulted in 1,173 DEGs at FDR<0.05 (1,846 DEGs at FDR<0.1) with a distribution of 61% (n=719 DEGs) up-regulated and 39% (n=454 DEGs) down-regulated genes (**Fig. 2E, F**; Supplementary). The top 10 dysregulated transcripts are mainly involved in the regulation of the innate immune system e.g., *Trim59*, and *Anln*, cell differentiation and survival e.g., *Ret*, *Disp3*, and *Mycl*, and the circadian rhythm e.g., *Hcrtr2*, *Ciart*, and *Npas4*. Also, among these highest significantly dysregulated genes, the MAP kinase phosphatases *Dusp4* and *Dusp6* were both significantly up-regulated in the alcohol drinking monkeys compared to alcohol-naive controls, which links the findings in monkeys to the MAPK-related outcomes identified in humans and rodents. STRING analysis did not yield any significantly interacting networks or genes on the protein level. GSEA resulted in a particular enrichment in cell signaling processes, such as “signal transduction”, “cell communication”, “cellular integrity”, “NCAM signaling for neurite out-growth”, and “response to stimulus”. Furthermore, MAPK-cascade regulatory processes, such as “MAPK signaling pathway”, “MAPK targets/ Nuclear events mediated by MAP kinases”, “ERK/MAPK targets”, and “AGE-RAGE signaling pathway in diabetic complication”, were highly over-represented, which further strengthens the importance of this pathway in alcohol dependent monkeys. Cell type enrichment analysis did not show any significant results.

### Cross-species comparison of the PFC results in DEGs that converge to an enrichment in neuroimmune system-related pathways, BBB regulation and MAPK-signaling

Since we were able to retrieve a sufficient number of datasets for the human, rodent, and monkey PFC, we next were interested in a direct comparison of the gene expression patterns across these species. Because we are expecting a higher degree of heterogeneity when comparing cross-species specific transcriptomic profiles, we set the threshold for DEGs to FDR<0.1. In the cross-species comparison, 13 common DEGs were identified (**Fig. 2G**). Among these, two genes were dysregulated in the same direction across all species, *AGBL4* and *TMEM80*. The remaining 11 transcripts were not dysregulated in the same direction across all three species (**Fig. 2G&H**). Furthermore, when comparing the direction of these 13 DEGs across the species in a pairwise manner, seven DEGs are dysregulated for humans and rodents, six for rodents and monkeys and four for humans and monkeys. The number of overlapping DEGs across the three species indicate that a higher number of DEGs in the human PFC are overlapping with the DEGs of the monkey PFC than with the rodent PFC (**Fig. 2G**). Cross-species specific STRING analyses identified *JUN* as a highly interconnected gene in humans and rodents. This gene has already been listed among the dysregulated DEGs with low FDR value in both of the species. However, in humans *JUN* is up-regulated (FDR(hum)=0.0004), while in the rodent PFC it is down-regulated (FDR(rod)=0.11). In monkeys, *JUN* is down-regulated, but does not reach significance (FDR(mon)=0.94). GSEA identified commonly dysregulated pathways in neuroimmune functions, vascular system regulation and BBB, cell cycle regulation, apoptosis and cancer-related pathways. Especially, the neuroimmune system-related pathways were impaired across humans, rodents and monkeys and the MAPK-related pathways. As the MAPK-pathways are also involved in the regulation of cell cycle and apoptosis, these hits from humans and monkeys also contributed to this category. The rodent GSEA mainly identified STAT-regulating mechanisms related to cell cycle-regulatory pathways, which were also present in the human down-stream analysis results. When further comparing the species-specific GSEA outcomes, in both humans as well as monkeys, mechanisms of BBB and regulation of vascular development and angiogenesis were significantly altered in the alcohol dependent phenotype.

### Meta-analysis of datasets derived from the NAc across human and rodent data results in a limited to absent number of DEGs

Systematic literature screening of the human postmortem brain studies resulted in three datasets derived from NAc tissue (n=241 samples), while for the rodent species, five datasets (n=77 samples) were identified and retrieved. Meta-analysis in the human NAc resulted in 17 DEGs at FDR<0.05 (44 DEGs at FDR<0.1), which were all up-regulated in AUD patients compared to healthy control individuals (**Fig. 3A, B**). The DEGs with the lowest FDR value were *ENPEP* (FDR=0.00035), *SERPINA3* (FDR=0.0042), *SLC7A2* (FDR=0.014), and *EDN1* (FDR=0.014). STRING analysis did not yield significantly inter-connected genes. GSEA resulted in dysregulated pathways of the neuroimmune system (e.g., “inflammatory response”, “response to cytokine”, “granulocyte migration”), blood vessel system (“leukocyte migration & chemotaxis”, “Fluid shear stress and atherosclerosis”, “blood vessel development”, “blood vessel morphogenesis”), cell integrity and signaling (“TNF signaling pathway”, “JAK-STAT signaling pathway”, “vesicle lumen”).

**Fig. 3:**
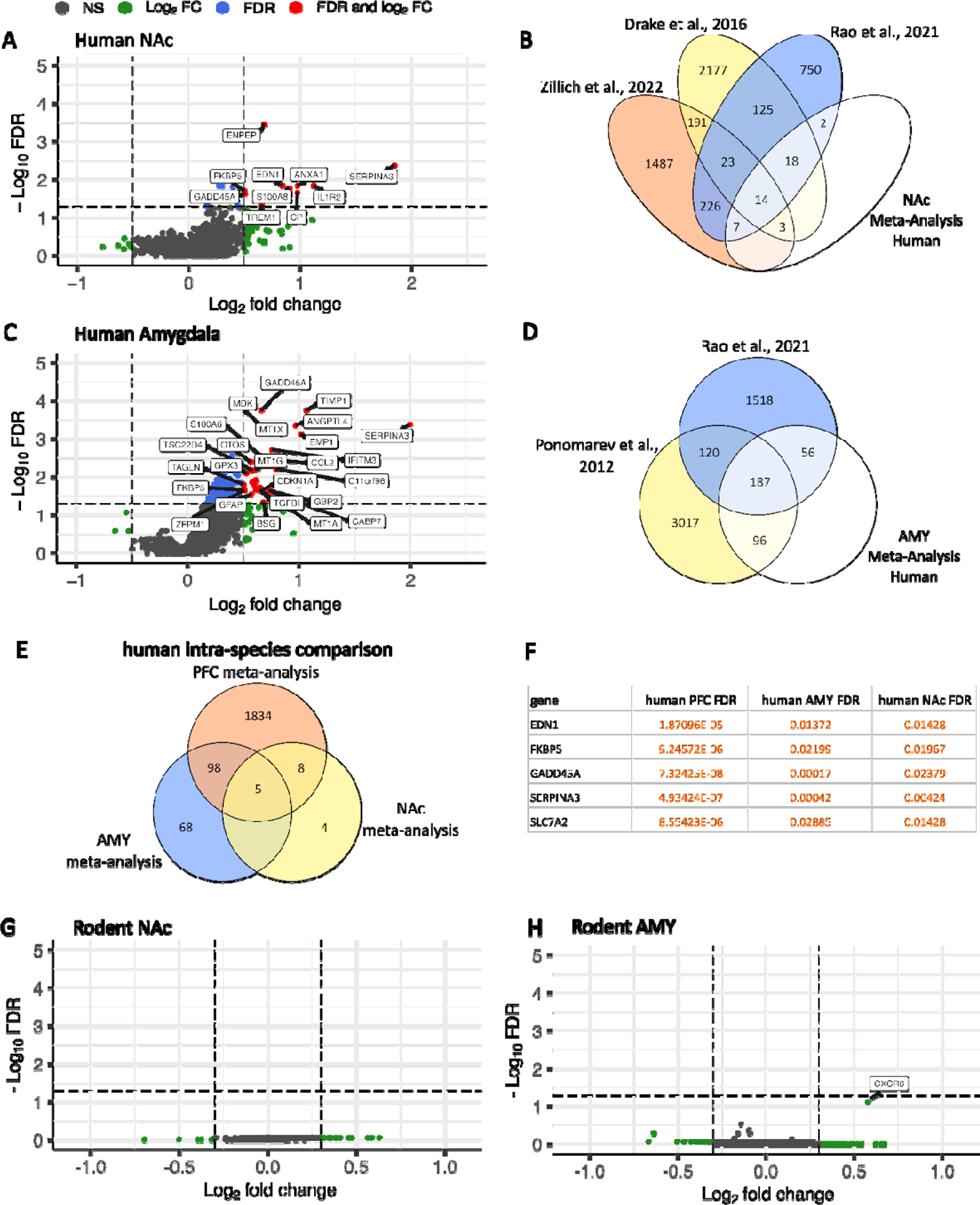
Meta-Analysis of the transcriptome-wide gene expression data from NAc and AMY from humans (A-F) and rodents (G, H) identified by Stouffer’s p-value combination with FDR<0.05 as threshold for significant DEGs. Transcripts with FDR<0.05 and log2 fold-change (FC)>0.5 are highlighted in red; DEGs with FDR<0.05 and FC<0.25 are highlighted in blue; DEGs with FDR>0.05 and FC>0.25 are highlighted in green; DEGs with FDR>0.05 and FC<0.5 for humans and FC<0.3 are highlighted in gray. (A) Gene expression pattern of the transcriptome-wide meta-analysis in human NAc. (B) Overlapping transcripts derived from human NAc original studied and the meta-analysis. (C) Gene expression pattern of the transcriptome-wide meta-analysis in human AMY. (D) Overlapping transcripts derived from human AMY original studied and the meta-analysis. (E) Venn diagram depicting the interspecies comparison considering the DEGs with FDR<0.05 across the human PFC, NAc and AMY meta-analysis results. (F) Five DEGs have been detected to be consistently up-regulated across these brain regions: *EDN*, *FKBP5*, *GADD45A*, *SERPINA3*, and *SLC7A2*. (G) Gene expression pattern of the transcriptome-wide meta-analysis in rodent NAc. (H) Gene expression pattern of the transcriptome-wide meta-analysis in rodent AMY. Considering a threshold of FDR<0.05, no DEGs were detected for both rodent NAc and AMY meta-analysis.

In the rodent analysis no DEGs at FDR<0.05 as well as FDR<0.1 appeared to be significant (**Fig. 3G**) and GSEA did not lead to any significant finding. When considering the mouse datasets separately, neither significant DEGs nor enriched pathways could be identified.

### Meta-analysis in the human AMY yielded DEGs that are mainly enriched in immune regulatory and BBB-related pathways, while in the rodent AMY no significant DEGs were detected

The meta-analysis of the AMY samples comprised six datasets from humans (n=92 samples) and two from rodents (n=88 samples). In the human meta-analysis, 171 DEGs with FDR<0.05 (339 DEGs with FDR<0.1) were identified with 100% of transcripts being up-regulated in AUD patients compared to healthy control individuals (**Fig. 3C, D**). Subsequent GSEA resulted in enrichment of these DEGs for dysregulated pathways in immune regulation (e.g., “inflammatory response”, “PPAR signaling pathway”, “IL-4 and IL-13 signaling”, “Cytokine signaling in immune system”), and factors of BBB integrity, such as “angiogenesis”, “hemostasis”, “blood vessels morphogenesis”, “regulation of vasculature development”, and “vasculature development”, and “JAK-STAT signaling pathway”. Pathways suggesting a direct influence on the protein level were also coming up, such as “negative regulation of hydrolase activity”, “regulation of endopeptidase activity”, and “post-translational protein phosphorylation”.

The rodent meta-analysis of the AMY included six datasets with a total of 88 animals. The gene expression analysis did not yield any significant DEGs at FDR<0.05 and only one DEG at FDR<0.1, namely *Cxcr6* (FDR=0.078) (**Fig. 3H**). GSEA did not result in any significant outcome for the rodent AMY. According to the sample descriptive that we extracted from the original studies, half of the datasets did not report the exact sub-region of the AMY. Since the AMY is known to be a very heterogeneous brain region that consists of multiple nuclei, we additionally analyzed the datasets that specifically stated the central AMY (CeA) (n(datasets)=3; n(animals)=43). These datasets contain mice as experimental animals and were conducted from the same research laboratory and the same first author (Supplementary). However, this analysis did not yield any significantly altered transcripts at FDR<0.05. At the threshold of FDR<0.1, one gene appeared as significant, which is *Cxcr6* (FDR=0.074) that already occurred in the rodent AMY meta-analysis after applying the same threshold for significance.

### Human cross-region analysis points towards new biomarkers for AUD diagnosis

As described in the sections above, humans were the only species that resulted in significantly altered transcripts within all the three brain regions analyzed in this study. Therefore, we wanted to determine whether there are transcripts that are dysregulated across these brain regions within the human species. As indicated in **Fig. 3E**, five DEGs are overlapping across PFC, NAc and AMY with FDR<0.05. These five transcripts are consistently up-regulated (**Fig. 3F**). We hypothesize that the DEGs that are dysregulated in the same direction across PFC, NAc and AMY, might be potential biomarkers for AUD, as they show a common pattern of dysregulation due to the alcohol dependent phenotype, irrespective of the brain region.

On the level of enriched pathways, there were common signatures observed across the PFC, NAc and AMY that point towards a general impairment of certain biological mechanisms. As already pointed out in the cross-species comparison of the PFC, also in the intra-species comparison the four main categories neuroimmunity, vascular system and BBB regulation, cell cycle regulation, apoptosis and cancer-related pathways such as MAPK signaling were dysregulated due to the AUD phenotype in all three brain regions with the majority of these terms being positively enriched. In addition, glia cell-specific terms, such as “regulation of glial cell apoptotic process”, “glia cell differentiation”, “gliogenesis” and “glial cell development” were enriched across the three brain regions.

## Discussion

The aim of this study was to conduct a comparative meta-analysis of transcriptome-wide data obtained from brain tissue samples collected from both animal models of alcohol dependence and individuals diagnosed with Alcohol Use Disorder (AUD) and healthy controls. Following a rigorous process of systematic literature screening and data collection, we acquired a sufficient number of datasets to perform meta-analyses for three specific brain regions: the PFC, the NAc, and the AMY. Key findings from this meta-analyzed 36 cross-species (mice, rats, monkeys, humans) transcriptome-wide RNA expression datasets with an alcohol-dependent phenotype, comprising a total of 1.000 samples, include: (i) the transcriptomic profiles of the PFC exhibit the most pronounced alterations across all three species and points out 13 common DEGs. (ii) The transcriptomic profile in the NAc appear to be less susceptible to long-term alcohol consumption followed by extended abstinence. (iii) Human intra-species comparison between the three brain areas suggests five potential biomarkers for AUD (e.g., SERPINA3). (iv) The most consistent finding across all meta-analyses performed here is the dysregulation of numerous genes encoding inflammatory processes and BBB integrity; in conjunction with the cell type-specific accumulation of these DEGs in astrocytes, microglia and endothelial cells, this finding is consistent with the proposed critical role of inflammatory mechanisms in AUD ^46–49^.

### Meta-analysis of the human and rodent PFC led to the majority of DEGs that are enriched in inflammatory processes

In the human PFC, 1,945 DEGs (FDR<0.05) resulted from the meta-analysis combining eight datasets with 360 samples. The volcano plot depicting the outcome DEGs of this analysis shows that the majority of DEGs is up-regulated. As a consequence, the majority of enriched pathways were positively enriched. The main finding is that a majority of DEGs is enriched in inflammatory processes. Previous research has repeatedly shown that especially inflammation - related genes are up-regulated in alcohol dependence ^46–48^. And furthermore, that neuroinflammatory processes indeed promote AUD development and progression ^49, 50^. Consequently, the role of anti-inflammatory approaches is suggested to be particularly promising as for AUD treatment ^51^. Furthermore, the cell type-specific enrichment in glia cells, such as astrocytes underlines the overrepresentation of inflammatory mechanisms in the PFC of AUD individuals. This finding is in line with one of the key hypotheses in the field, namely that glial cells, such as astrocytes and microglia, are considered as new cellular targets for treating alcohol-induced inflammatory and behavioral responses (Crews et al., 2017; 2023; Erickson et al., 2019; Grantham et al., 2023).

The human PFC meta-analysis results are in line with the findings in rodents. Thus, rodent GSEA also pointed towards a major impairment of immune system-related pathways, such as leukocyte functioning, cytokine production, and especially T cell regulatory mechanisms. Indeed, T cell functioning seems to be altered by chronic alcohol consumption in both AUD patients and rodent models thereof. Thus, a recent methylation study on CD3+ T cells of AUD patients and matched controls identified numerous methylation sites to be altered in the AUD condition ^52^, while epigenetic (chromatin immunoprecipitation [ChIP]-seq) analysis in the PFC of post-dependent rats showed significant enrichment in the interleukin signaling pathway resulting in a diminished anti-inflammatory IL-6 response ^22, 53^.

### DEGs in the human PFC that encode for the metallothionein (MT) family may explain zinc deficiency in AUD patients

The DEG of the human PFC meta-analysis with the lowest FDR (FDR=1.5E-10), *MT1X*, is a gene that encodes for metallothionein 1X. Additional members of the metallothionein (MT) family, such as *MT1A*, *MT1M*, *MT1E*, *MT1G*, *MT2A* and *MT3* reached significance in our study, as well, and all of these were up-regulated in the AUD patient samples. MT genes are involved in zinc ion binding activity and zinc, in turn, is a key element for the activation and binding of certain transcription factors through its participation in the zinc finger region of the protein. MT genes have been associated with AUD in previous studies. MT genes were found to be elevated in the hippocampus of AUD patients ^54^. Another study that focused on mRNA and miRNA transcription patterns in the NAc and PFC of AUD patients and matched controls, identified similar MT gene candidates (*MT1E*, *MT1F*, *MT1G*, *MT1H*, *MT1HL1*, *MT1X*, *MT2A*, and *MT3*) being up-regulated in both brain regions investigated ^55^. The upregulation of MT genes explains the well-known phenomenon of zinc deficiency in AUD a phenomenon that may contribute to disease maintenance ^56, 57^. Because of zinc deficiency, a number of zinc finger proteins are known to be impaired by AUD, which can be observed in our data, as well. In sum, it is concluded that not only daily oral zinc supplementation but also regulation of zinc homeostasis through MTs might be helpful for treating AUD patients.

### Meta-analysis of the monkey PFC results in alteration of MAPK/ERK-signaling mainly by *Dusp4*

The meta-analysis of the monkey PFC datasets resulted in 1,173 DEGs (FDR<0.05). The DEG of the monkey PFC meta-analysis with the lowest FDR (FDR=6.1E-24 and FC 1.3), *Dusp4*, is a gene that encodes for the Dual-Specificity Protein Phosphatase 4 which is a major inhibitor of the MAPK pathway and is also involved in circadian rhythmicity ^58–61^. So far, in alcohol-related research it was reported that *Dusp4* is involved in ethanol-induced liver injury, where an increase of *Dusp4* expression was suggested to cause autophagy via miR-26a ^62^. In the monkey PFC results, *Dusp4* was not the only family member of DUSPs that were significantly altered in the chronic alcohol-drinking monkeys. In total, seven DUSPs were identified as dysregulated with five up-regulated (*Dusp1*, *Dusp14*, *Dusp4*, *Dusp5*, *Dusp6*) transcripts. As one of the major interaction partners of MAPK/ERK pathway-related components, it does not come with a surprise that MAPK- and ERK-signaling is the most frequently represented enriched pathway in the GSEA. Additionally, five MAPK genes were dysregulated in the monkey PFC, which further strengthens the impairment of this signaling pathway. *Dusp4* regulates several components of the MAPK pathways and thereby also impacts on circadian control ^58^. In this respect, it is of note that circadian rhythmicity is severely altered in AUD patients and in animal models of alcohol dependence ^63^ and *Dusp4* should thus be considered as a new target to normalize circadian rhythmicity in AUD patients.

In conclusion, the monkey meta-analyzed data are in line with numerous publications showing the importance of the MAPK/ERK pathway in mediating acute and long-term effects of alcohol ^64–67^. However, this is not an alcohol specific effect since reinforcing effects of other drugs of abuse, such as cocaine ^68, 69^ and nicotine ^70, 71^ are also mediated by this pathway. Therefore, clinically relevant, systemically administered drugs that target ERK signaling in the brain could be used to treat AUD and other substance use disorders ^68^.

### Convergent transcriptomic signature in the PFC across AUD patients and alcohol-dependent rodents and monkeys

The inter-species comparison resulted in 13 DEGs that were significant across all three species with two genes - *AGBL4* and *TMEM80* - being dysregulated in the same direction. ATP/GTP binding protein-like 4 (AGBL4) was found to be associated with alcohol dependence ^72^ and smoking ^73^ in recent GWAS studies; and smoking is the most commonly co-occurring psychiatric comorbidity in AUD patients. The function and relevance of transmembrane protein 80 (*TMEM80*) in health and disease is currently not known. However, another DEG that encodes for transmembrane protein *TMEM229A* also emerged in all three species and a particular gene variant of *TMEM229A* is associated with alcohol consumption in a GWAS ^74^. Another DEG with convergent transcriptomic signature for all three species is *HCRTR2* that encodes for the orexin receptor 2. The involvement of orexin receptor 2 in alcohol consumption ^75^ and relapse-like behavior in alcohol dependent rats ^76–78^ supports our convergent cross-species finding.

However, the focus on single gene findings in a disease with high heterogeneity such as AUD might not be the best way for medication development. Hence, the investigation of dysregulated pathways and physiological processes may lead to more promising targets for intervention in AUD patients. In this respect, the dysregulation of the integrity and function of the BBB stands out in our convergent analysis. Since the BBB is regulating the in- and efflux of components derived from the periphery, it protects the brain from inflammatory factors derived from the periphery. Our cell type enrichment analysis resulted in endothelial cells as the strongest enriched cell type in the PFC. Since endothelial cells are one of the principal components making up the BBB, DEGs within endothelial cells of the BBB may contribute to a physiological dysfunction of BBB integrity and will thereby contribute to the aforementioned inflammatory processes in the PFC and other brain sites. The impairment of the BBB due endothelial cell dysfunction and subsequent inflammatory brain responses have been reported previously and re-establishing the BBB integrity has been suggested as a promising therapeutic tool in abstinence and withdrawal ^79–82^.

Consistent with the result of the convergent analysis in the PFC, we also found numerous DEGs in the meta-analysis of the human AMY that were mainly enriched in inflammatory processes and BBB integrity, supporting the idea of a general pathological mechanism. The chain of pathological processes can include the toxic effect of alcohol, which leads to peripheral inflammatory processes and injuries to various organs. Thus, the toxic effect of alcohol may cause blood-brain barrier leakage due to damage to the endothelial cells, allowing mediators of inflammation to enter the brain. Some brain areas, particularly the PFC, may be more sensitive to these processes and then suffer cell death and loss of function.

### The transcriptomic profile of the NAc seems to be less vulnerable to long-term alcohol consumption followed by protracted abstinence

The NAc, also known as the center of reward processing, has an essential role in the development of AUD mainly, where alcohol is experienced as pleasurable and rewarding ^83–85^. Our meta-analysis of human and rodent NAc transcriptomic profiles showed limited to no significant findings in alcohol dependent subjects compared to non-dependent controls. Since this study is focused on more advanced stages of AUD, it could well be that the NAc transcriptome is less involved during the maintenance of the alcohol dependence.

Previous research describes that the NAc may mostly be involved in the initiation of AUD characterized by the binge and acquisition phase ^86, 87^. The data that we included in our analyses also underline these findings, as there are no DEGs reported in the rodent studies and very little in the human postmortem studies. A recent single nucleus RNA sequencing (snRNA-Seq) on the NAc of human postmortem brain ^88^ reported 26 DEGs that were mainly attributed to D1- and D2 medium spiny neurons, oligodendrocytes, and microglia. In comparison, another snRNA-Seq study that was performed on human postmortem PFC identified 10-fold more DEGs (253 DEGs at FDR<0.05) ^89^. Therefore, our study supports the notion that the NAc has only little relevance in the maintenance of AUD. Alternatively, the lack of transcriptomic response could reflect a form of alcohol-induced anaplasticity. Such an inability to adaptively respond to specific challenges or stimuli has been observed in some animal models of addiction ^90^ as well as certain tumor cells ^91^.

### Human intra-species comparison suggests five potential biomarkers for AUD

Since significant DEGs for all three brain regions were only observed in AUD patients, an intra-species comparison was performed to identify potential commonly dysregulated genes in these brain regions. Identified genes or their combinations may represent potential biomarkers for AUD that could lead to the much-needed improvement in the diagnosis of AUD. Five DEGs were overlapping across PFC, NAc and AMY – namely *EDN1*, *FKBP5*, *GADD45A*, *SLC7A2*, and *SERPINA3* – and all of these genes were consistently up-regulated. For example, *SERPINA3* might be a promising target for biomarker identification, as previous research on hippocampus and peripheral blood of AUD patients reported up-regulated *SERPINA3* gene expression, as well ^54, 92^. However, upregulation of *SERPIN3A* also occurs in the aging brain and in neurodegenerative diseases ^93^ and is therefore not only specific to AUD disease. In any case, it seems illusory that a psychiatric diagnosis can be based on a single gene change; rather, the combination of several genetic markers might be more reasonable.

## Conclusion

The meta-analyzed, transcriptome-wide, cross-species data provided here represents a compendium of genes, signaling pathways, and physiological and cellular processes that are altered in AUD. Future studies should focus on functional validation of the DEGs reported here, particularly those that converge across all three species studied. The pathomechanisms described here, which include damage to BBB integrity by endothelial cell-enriched DEGs and subsequent inflammatory processes in vulnerable brain regions such as PFC and AMY, warrant further research, and drugs that counteract these pathomechanisms may indeed be useful future treatments for AUD. Furthermore, our results once again underline the notion that the NAc is not of great interest for studies on addictive processes, while there is no doubt that the NAc plays a crucial role in the initiation and acquisition of alcohol drinking behavior. Finally, the combination of DEGs proposed here, which are commonly upregulated in brain tissue and probably also in the periphery, should be further investigated in the context of the biological diagnosis of AUD.

## Supporting information

Summary Statistics Analysis Human

Summary Statistics Analysis Monkeys

Summary Statistics Analysis Rodents

Supporting general informations

## Acknowledgements

The authors thank the CAMARADES research group Edinburgh for their very helpful support on the study design and the systematic literature screening and data extraction process. In addition, we are very grateful to all the corresponding authors of the original studies that made it possible to include these datasets in our analyses. Without this involvement, it would not have been possible to conduct the study to this extent. The authors declare no biomedical financial interests or potential conflict of interest.

## Author contribution

MMF, AMB, FG and RS designed the study. MMF, AMB and FG designed the keywords for systematic literature screening. MMF, ECT, AMB and FG performed systematic literature screening and data extraction. MMF and WHS contacted the corresponding authors in case datasets were not publicly available. MMF, ECT, MJWH and RS designed the analysis plan. MMF, ECT and MJWH performed data analysis. ACH, MAP, WHS, SF and RH provided unpublished data. MAP, JS, WHS, RDM, and RS provided scientific support. MMF and RS wrote the manuscript. All authors have read and approved the manuscript.

## Data Availability Statement

The meta-analysis in rodents (Open Science Framework: https://doi.org/10.17605/OSF.IO/TF8R4) as well as in human post-mortem brain tissue (Prospero: CRD42020192453) has been pre-registered in advance. As mentioned in the methods section, the analysis strategy was deviating from the pre-registered methods, since the obtained data were not enabling the initially planned random effect size combination. Datasets as well as analysis pipelines will be publicly available from the day of publication onwards.

## References

1. World Health Organization W. Global status report on alcohol and health 2018. Geneva: World Health Organization; 20182018.

2. MacKillop J, Agabio R, Feldstein Ewing SW, Heilig M, Kelly JF, Leggio L, et al. Hazardous drinking and alcohol use disorders. Nat Rev Dis Primers 2022; 8(1): 80.

3. GBD. Alcohol use and burden for 195 countries and territories, 1990-2016: a systematic analysis for the Global Burden of Disease Study 2016. Lancet 2018; 392(10152): 1015–1035.

4. Heilig M, MacKillop J, Martinez D, Rehm J, Leggio L, Vanderschuren LJMJ. Addiction as a brani disease revised: why it still matters, and. the need for consilience. Neuropharmacology 2021; 46(10): 1715–1723.

5. Heilig M, Augier E, Pfarr S, Sommer WH. Developing neuroscience-based treatments for alcohol addiction: A matter of choice? Transl Psychiatry 2019; 9(1): 255.

6. Burnette EM, Nieto SJ, Grodin EN, Meredith LR, Hurley B, Miotto K et al. Novel Agents for the Pharmacological Treatment of Alcohol Use Disorder. Drugs 2022; 82(3): 251–274.

7. Kranzler HR, Soyka M. Diagnosis and Pharmacotherapy of Alcohol Use Disorder: A Review. JAMA 2018; 320(8): 815–824.

8. Witkiewitz K, Litten RZ, Leggio L. Advances in the science and treatment of alcohol use disorder. Sci Adv 2019; 5(9): eaax4043.

9. Spanagel R. Alcoholism: a systems approach from molecular physiology to addictive behavior. Physiol Rev 2009; 89(2): 649–705.

10. Ron D, Barak S. Molecular mechanisms underlying alcohol-drinking behaviours. Nat Rev Neurosci 2016; 17(9): 576–591.

11. Abrahao KP, Salina AG, Lovinger DM. Alcohol and the Brain: Neuronal Molecular Targets, Synapses, and Circuits. Neuron 2017; 96(6): 1223–1238.

12. Egervari G, Siciliano CA, Whiteley EL, Ron D. Alcohol and the brain: from genes to circuits. Trends Neurosci 2021; 44(12): 1004–1015.

13. Koob GF, Volkow ND. Neurocircuitry of addiction. Neuropsychopharmacology 2010; 35(1): 217–238.

14. Noori HR, Spanagel R, Hansson AC. Neurocircuitry for modeling drug effects. Addict Biol 2012; 17(5): 827–864.

15. Button KS, Ioannidis J, Mokrysz C, Nosek BA, Flint J, Robinson ES et al. Power failure: why small sample size undermines the reliability of neuroscience. Nature Reviews Neuroscience 2013; 14(5): 365–376.

16. Spanagel R. Ten Points to Improve Reproducibility and Translation of Animal Research. Front Behav Neurosci 2022; 16: 869511.

17. Farris SP, Arsappan D, Hunicke-Smith S, Harris RA, Mayfield RD. Transcriptome organization for chronic alcohol abuse in human brain. Mol Psychiatry 2015; 20(11): 1438–1447.

18. Ponomarev I, Wang S, Zhang L, Harris RA, Mayfield RD. Gene coexpression networks in human brain identify epigenetic modifications in alcohol dependence. J Neurosci 2012; 32(5): 1884–1897.

19. Rao X, Thapa KS, Chen AB, Lin H, Gao H, Reiter JL et al. Allele-specific expression and high-throughput reporter assay reveal functional genetic variants associated with alcohol use disorders. Mol Psychiatry 2021; 26(4): 1142–1151.

20. Zillich L, Poisel E, Frank J, Foo JC, Friske MM, Streit F et al. Multi-omics signatures of alcohol use disorder in the dorsal and ventral striatum. Transl Psychiatry 2022; 12(1): 190.

21. Meinhardt MW, Sommer WH. Postdependent state in rats as a model for medication development in alcoholism. Addict Biol 2015; 20(1): 1–21.

22. Heilig M, Barbier E, Johnstone AL, Tapocik J, Meinhardt MW, Pfarr S et al. Reprogramming of mPFC transcriptome and function in alcohol dependence. Genes Brain Behav 2017; 16(1): 86–100.

23. Kapoor M, Wang JC, Farris SP, Liu Y, McClintick J, Gupta I et al. Analysis of whole genome-transcriptomic organization in brain to identify genes associated with alcoholism. Transl Psychiatry 2019; 9(1): 89.

24. Liu J, Lewohl JM, Harris RA, Iyer VR, Dodd PR, Randall PK et al. Patterns of gene expression in the frontal cortex discriminate alcoholic from nonalcoholic individuals. Neuropsychopharmacology 2006; 31(7): 1574–1582.

25. Manzardo AM, Gunewardena S, Butler MG. Over-expression of the miRNA cluster at chromosome 14q32 in the alcoholic brain correlates with suppression of predicted target mRNA required for oligodendrocyte proliferation. Gene 2013; 526(2): 356–363.

26. Wang F, Gelernter J, Zhang H. Differential Expression of miR-130a in Postmortem Prefrontal Cortex of Subjects with Alcohol Use Disorders. J Addict Res Ther 2013; 4(155).

27. Drake J, McMichael GO, Vornholt ES, Cresswell K, Williamson V, Chatzinakos C et al. Assessing the Role of Long Noncoding RNA in Nucleus Accumbens in Subjects With Alcohol Dependence. Alcohol Clin Exp Res 2020; 44(12): 2468–2480.

28. Hade AC, Philips MA, Reimann E, Jagomae T, Eskla KL, Traks T et al. Chronic Alcohol Use Induces Molecular Genetic Changes in the Dorsomedial Thalamus of People with Alcohol-Related Disorders. Brain Sci 2021; 11(4).

29. Meinhardt MW, Hansson AC, Perreau-Lenz S, Bauder-Wenz, Staehlin O, Heilig M et al. Rescue of Infralimbic mGluR2 Deficit Restores Control Over Drug-Seeking Behavior in Alcohol Dependence. Journal of Neuroscience 2013; 33(7): 2794–2806.

30. Osterndorff-Kahanek EA, Becker HC, Lopez MF, Farris SP, Tiwari GR, Nunez YO et al. Chronic Ethanol Expsoure Produces Time- and Brain Region-Dependent Changes in Gene Coexpression Networks. PLoS One 2015; 10(3): e0121522.

31. Smith ML, Lopez MF, Archer KJ, Wolen AR, Becker HC, Miles MF. Time-Course Analysis of Brain Regional Expression Network Responses to Chronic Intermittent Ethanol and Withdrawal: Implications for Mechanisms Underlying Excessive Ethanol Consumption. PLoS One 2016; 11(1): e0146257.

32. Smith ML, Lopez MF, R. WA, Becker HC, Miles MF. Brain regional gene expression network analysis identifies unique interactions between chronic ethanol exposure and consumption. PLoS One 2020; 10(3): e0121522.

33. Farris SP, Tiwari GR, Ponomareva O, Lopez MF, Mayfield RD, Becker HC. Transcriptome Analysis of Alcohol Drinking in Non-Dependent and Dependent Mice Following Repeated Cycles of Forced Swim Stress Exposure. Brain Sci 2020; 10(5).

34. Bogenpohl JW, Smith ML, Farris SP, Dumur CI, Lopez MF, Becker HC et al. Cross-Species Co-analysis of Prefrontal Cortex Chronic Ethanol Transcriptome Responses in Mice and Monkeys. Front Mol Neurosci 2019; 12: 197.

35. Walter NAR, Zheng CL, Searles RP, McWeeney SK, Grant KA, Hitzemann R. Chronic Voluntary Ethanol Drinking in Cynomolgus Macaques Elicits Gene Expression Changes in Prefrontal Cortical Area 46. Alcohol Clin Exp Res 2020; 44(2): 470–478.

36. Burguillo FJ, Martin J, Barrera I, Bardsley WG. Meta-analysis of microarray data: The case of imatinib resistance in chronic myelogenous leukemia. Comput Biol Chem 2010; 34(3): 184–192.

37. Panahi B, Frahadian M, Dums JT, Hejazi MA. Integration of Cross Species RNA-seq Meta-Analysis and Machine-Learning Models Identifies the Most Important Salt Stress-Responsive Pathways in Microalga Dunaliella. Front Genet 2019; 10: 752.

38. Rau A, Marot G, Jaffrezic F. Differential meta-analysis of RNA-seq data from multiple studies. BMC Bioinformatics 2014; 15: 91.

39. Stouffer SA, Suchman EA, DeVinney LC, Star SA, Williams Jr. RM. Studies in Social Psychology in World War II: The American Soldier. Vol. 1, Adjustment During Army Life. Princeton: Princeton University Press 1949.

40. Marot G, Foulley JL, Mayer CD, Jaffrezic F. Moderated effect size and P-value combinations for microarray meta-analyses. Bioinformatics 2009; 25(20): 2692–2699.

41. Ryu SY, Wendt GA. MetaMSD: meta analysis for mass spectrometry data. PeerJ 2019; 7: e6699.

42. Zaykin DV. Optimally weighted Z-test is a powerful method for combining probabilities in meta-analysis. J Evol Biol 2011; 24(8): 1836–1841.

43. Wu T, Hu E, Xu S, Chen M, Guo P, Dai Z et al. clusterProfiler 4.0: A universal enrichment tool for interpreting omics data. The Innovation 2021; 2(3): 100141.

44. Raudvere U, Kolberg L, Kuzmin I, Arak T, Adler P, Peterson H et al. g:Profiler: a web server for functional enrichment analysis and conversions of gene lists (2019 update). Nucleic Acids Res 2019; 47(W1): W191–W198.

45. Ma W, Sharma S, Jin P, Gourley SL, Qin ZS. LRcell: detecting the source of differential expression at the sub-cell-type level from bulk RNA-seq data. Brief Bioinform 2022; 23(3).

46. Crews FT, Lawrimore CJ, Walter TJ, Coleman LG, Jr. The role of neuroimmune signaling in alcoholism. Neuropharmacology 2017; 122: 56–73.

47. Erickson EK, Blednov YA, Harris RA, Mayfield RD. Glial gene networks associated with alcohol dependence. Sci Rep 2019; 9(1): 10949.

48. Grantham EK, Barchiesi R, Salem NA, Mayfield RD. Neuroimmune pathways as targets to reduce alcohol consumption. Pharmacol Biochem Behav 2023; 222: 173491.

49. De Santis S, Cosa-Linan A, Garcia-Hernandez R, Dmytrenko L, Vargova L, Vorisek I et al. Chronic alcohol consumption alters extracellular space geometry and transmitter diffusion in the brain. Sci Adv 2020; 6(26): eaba0154.

50. Perez-Cervera L, De Santis S, Marcos E, Ghorbanzad-Ghaziany Z, Trouve-Carpena A, Selim MK et al. Alcohol-induced damage to the fimbria/fornix reduces hippocampal-prefrontal cortex connection during early abstinence. Acta Neuropathol Commun 2023; 11(1): 101.

51. Cruz B, Vozella V, Carper BA, Xu JC, Kirson D, Hirsch S et al. FKBP5 inhibitors modulate alcohol drinking and trauma-related behaviors in a model of comorbid post-traumatic stress and alcohol use disorder. Neuropsychopharmacology 2023; 48(8): 1144–1154.

52. Bruckmann C, Islam SA, MacIsaac JL, Morin AM, Karle KN, Di Santo A et al. DNA methylation signatures of chronic alcohol dependence in purified CD3(+) T-cells of patients undergoing alcohol treatment. Sci Rep 2017; 7(1): 6605.

53. Johnstone AL, Andrade NS, Barbier E, Khomtchouk BB, Rienas CA, Lowe K et al. Dysregulation of the histone demethylase KDM6B in alcohol dependence is associated with epigenetic regulation of inflammatory signaling pathways. Addict Biol 2021; 26(1): e12816.

54. McClintick JN, Xuei X, Tischfield JA, Goate A, Foroud T, Wetherill L et al. Stress-response pathways are altered in the hippocampus of chronic alcoholics. Alcohol 2013; 47(7): 505–515.

55. Vornholt E, Drake J, Mamdani M, McMichael G, Taylor ZN, Bacanu SA et al. Network preservation reveals shared and unique biological processes associated with chronic alcohol abuse in NAc and PFC. PLoS One 2020; 15(12): e0243857.

56. Morud J, Adermark L, Ericson M, Soderpalm B. Alterations in ethanol-induced accumbal transmission after acute and long-term zinc depletion. Addict Biol 2015; 20(1): 170–181.

57. Skalny AV, Skalnaya MG, Grabeklis AR, Skalnaya AA, Tinkov AA. Zinc deficiency as a mediator of toxic effects of alcohol abuse. Eur J Nutr 2018; 57(7): 2313–2322.

58. Hamnett R, Crosby P, Chesham JE, Hastings MH. Vasoactive intestinal peptide controls the suprachiasmatic circadian clock network via ERK1/2 and DUSP4 signalling. Nat Commun 2019; 10(1): 542.

59. Kirchner A, Bagla S, Dachet F, Loeb JA. DUSP4 appears to be a highly localized endogenous inhibitor of epileptic signaling in human neocortex. Neurobiol Dis 2020; 145: 105073.

60. Ephrame SJ, Cork GK, Marshall V, Johnston MA, Shawa J, Alghusen I et al. O-GlcNAcylation regulates extracellular signal-regulated kinase (ERK) activation in Alzheimer’s disease. Front Aging Neurosci 2023; 15: 1155630.

61. Prabhakar S, Asuthkar S, Lee W, Chigurupati S, Zakharian E, Tsung AJ et al. Targeting DUSPs in glioblastomas - wielding a double-edged sword? Cell Biol Int 2014; 38(2): 145–153.

62. Han W, Fu X, Xie J, Meng Z, Gu Y, Wang X et al. MiR-26a enhances autophagy to protect against ethanol-induced acute liver injury. J Mol Med (Berl) 2015; 93(9): 1045–1055.

63. Spanagel R, Rosenwasser AM, Schumann G, Sarkar DK. Alcohol consumption and the body’s biological clock. Alcohol Clin Exp Res 2005; 29(8): 1550–1557.

64. Sanna PP, Simpson C, Lutjens R, Koob G. ERK regulation in chronic ethanol exposure and withdrawal. Brain Res 2002; 948(1-2): 186–191.

65. Agoglia AE, Sharko AC, Psilos KE, Holstein SE, Reid GT, Hodge CW. Alcohol alters the activation of ERK1/2, a functional regulator of binge alcohol drinking in adult C57BL/6J mice. Alcohol Clin Exp Res 2015; 39(3): 463–475.

66. Faccidomo S, Salling MC, Galunas C, Hodge CW. Operant ethanol self-administration increases extracellular-signal regulated protein kinase (ERK) phosphorylation in reward-related brain regions: selective regulation of positive reinforcement in the prefrontal cortex of C57BL/6J mice. Psychopharmacology (Berl) 2015; 232(18): 3417–3430.

67. Hansson AC, Rimondini R, Neznanova O, Sommer WH, Heilig M. Neuroplasticity in brain reward circuitry following a history of ethanol dependence. Eur J Neurosci 2008; 27(8): 1912–1922.

68. Papale A, Morella IM, Indrigo MT, Bernardi RE, Marrone L, Marchisella F et al. Impairment of cocaine-mediated behaviours in mice by clinically relevant Ras-ERK inhibitors. Elife 2016; 5.

69. Bernardi RE, Olevska A, Morella I, Fasano S, Santos E, Brambilla R et al. The Inhibition of RasGRF2, But Not RasGRF1, Alters Cocaine Reward in Mice. J Neurosci 2019; 39(32): 6325–6338.

70. Valjent E, Pages C, Herve D, Girault JA, Caboche J. Addictive and non-addictive drugs induce distinct and specific patterns of ERK activation in mouse brain. Eur J Neurosci 2004; 19(7): 1826–1836.

71. Morella I, Pohorala V, Calpe-Lopez C, Brambilla R, Spanagel R, Bernardi RE. Nicotine self-administration and ERK signaling are altered in RasGRF2 knockout mice. Front Pharmacol 2022; 13: 986566.

72. Sun Y, Chang S, Liu Z, Zhang L, Wang F, Yue W et al. Identification of novel risk loci with shared effects on alcoholism, heroin, and methamphetamine dependence. Mol Psychiatry 2021; 26(4): 1152–1161.

73. Xu K, Li B, McGinnis KA, Vickers-Smith R, Dao C, Sun N et al. Genome-wide association study of smoking trajectory and meta-analysis of smoking status in 842,000 individuals. Nat Commun 2020; 11(1): 5302.

74. Adkins DE, Clark SL, Copeland WE, Kennedy M, Conway K, Angold A et al. Genome-Wide Meta-Analysis of Longitudinal Alcohol Consumption Across Youth and Early Adulthood. Twin Res Hum Genet 2015; 18(4): 335–347.

75. Brown RM, Khoo SY, Lawrence AJ. Central orexin (hypocretin) 2 receptor antagonism reduces ethanol self-administration, but not cue-conditioned ethanol-seeking, in ethanol-preferring rats. Int J Neuropsychopharmacol 2013; 16(9): 2067–2079.

76. Amodeo LR, Wills DN, Sanchez-Alavez M, Ehlers CL. Effects of an Orexin-2 Receptor Antagonist on Sleep and Event-Related Oscillations in Female Rats Exposed to Chronic Intermittent Ethanol During Adolescence. Alcohol Clin Exp Res 2020; 44(7): 1378–1388.

77. Flores-Ramirez FJ, Varodayan FP, Patel RR, Illenberger JM, Di Ottavio F, Roberto M et al. Blockade of orexin receptors in the infralimbic cortex prevents stress-induced reinstatement of alcohol-seeking behaviour in alcohol-dependent rats. Br J Pharmacol 2023; 180(11): 1500–1515.

78. Aldridge GM, Zarin TA, Brandner AJ, George O, Gilpin NW, Repunte-Canonigo V et al. Effects of single and dual hypocretin-receptor blockade or knockdown of hypocretin projections to the central amygdala on alcohol drinking in dependent male rats. Addict Neurosci 2022; 3.

79. Carrino D, Branca JJV, Becatti M, Paternostro F, Morucci G, Gulisano M et al. Alcohol-Induced Blood-Brain Barrier Impairment: An In Vitro Study. Int J Environ Res Public Health 2021; 18(5).

80. Rubio-Araiz A, Porcu F, Perez-Hernandez M, Garcia-Gutierrez MS, Aracil-Fernandez MA, Gutierrez-Lopez MD et al. Disruption of blood-brain barrier integrity in postmortem alcoholic brain: preclinical evidence of TLR4 involvement from a binge-like drinking model. Addict Biol 2017; 22(4): 1103–1116.

81. Somkuwar SS, Fannon MJ, Bao Nguyen T, Mandyam CD. Hyper-oligodendrogenesis at the vascular niche and reduced blood-brain barrier integrity in the prefrontal cortex during protracted abstinence. Neuroscience 2017; 362: 265–271.

82. Wei J, Dai Y, Wen W, Li J, Ye LL, Xu S et al. Blood-brain barrier integrity is the primary target of alcohol abuse. Chem Biol Interact 2021; 337: 109400.

83. Ho AL, Salib AN, Pendharkar AV, Sussman ES, Giardino WJ, Halpern CH. The nucleus accumbens and alcoholism: a target for deep brain stimulation. Neurosurg Focus 2018; 45(2): E12.

84. Scofield MD, Heinsbroek JA, Gipson CD, Kupchik YM, Spencer S, Smith AC et al. The Nucleus Accumbens: Mechanisms of Addiction across Drug Classes Reflect the Importance of Glutamate Homeostasis. Pharmacol Rev 2016; 68(3): 816–871.

85. Spiga S, Talani G, Mulas G, Licheri V, Fois GR, Muggironi G et al. Hampered long-term depression and thin spine loss in the nucleus accumbens of ethanol-dependent rats. Proc Natl Acad Sci U S A 2014; 111(35): E3745–3754.

86. Koob GF, Le Moal M. Plasticity of reward neurocircuitry and the ‘dark side’ of drug addiction. Nat Neurosci 2005; 8(11): 1442–1444.

87. Korber C, Sommer WH. From ensembles to meta-ensembles: Specific reward encoding by correlated network activity. Front Behav Neurosci 2022; 16: 977474.

88. van den Oord E, Xie LY, Zhao M, Aberg KA, Clark SL. A single-nucleus transcriptomics study of alcohol use disorder in the nucleus accumbens. Addict Biol 2023; 28(1): e13250.

89. Brenner E, Tiwari GR, Kapoor M, Liu Y, Brock A, Mayfield RD. Single cell transcriptome profiling of the human alcohol-dependent brain. Hum Mol Genet 2020; 29(7): 1144–1153.

90. Kasanetz F, Deroche-Gamonet V, Berson N, Balado E, Lafourcade M, Manzoni O et al. Transition to addiction is associated with a persistent impairment in synaptic plasticity. Science 2010; 328(5986): 1709–1712.

91. Khan SU, Khan MU, Khan MI, Fadahunsi AA, Khan A, Gao S et al. Role of circular RNAs in disease progression and diagnosis of cancers: An overview of recent advanced insights. Int J Biol Macromol 2022; 220: 973–984.

92. Zhang J, Powell CA, Kay MK, Sonkar R, Meruvu S, Choudhury M. Effect of Chronic Western Diets on Non-Alcoholic Fatty Liver of Male Mice Modifying the PPAR-γ Pathway via miR-27b-5p Regulation. Int J Mol Sci 2021; 22(4).

93. Vanni S, Colini Baldeschi A, Zattoni M, Legname G. Brain aging: A Ianus-faced player between health and neurodegeneration. J Neurosci Res 2020; 98(2): 299–311.

