## Supporting general informations for "A Systematic Review and Meta-analysis on the Transcriptomic Signatures in Alcohol Use Disorder"

**Supplementary Material**

**Suppl. Tab. 1**: Detailed information about included datasets and respective studies for meta-analysis in rodents.

| *Study* | *n* | *Strain* | *Treatment* | *Exposure length* | *Abstinence length* | *Brain Region* | *Method* |
| --- | --- | --- | --- | --- | --- | --- | --- |
| Meinhardt et al., 2013 | 18 (PFC)  16 (Amy)  14 (NAc) | Wistar Rats | CIE Vapor | 7 wk | 3 wk | mPFC,  Amy,  NAc | Microarray |
| Meinhardt et al., 2013 | 16 (PFC)  13 (Amy) | Wistar Rats | CIE Vapor | 4 wk | 3 wk | mPFC,  Amy | Microarray |
| Osterndorff-Kahanek et al., 2015 | 16 (PFC)  16 (Amy)  16 (NAc) | C57BL/6 Mice | CIE Vapor | 8 wk | 120 h | PFC,  Amy,  NAc | Microarray |
| Smith et al., 2016 | 12 (PFC)  12 (CeA)  11 (NAc) | C57BL/6 Mice | CIE Vapor | 6 wk | 72 h | PFC,  CeA,  NAc | Microarray |
| Smith et al., 2016 | 12 (PFC)  12 (CeA)  12 (NAc) | C57BL/6 Mice | CIE Vapor | 6 wk | 7 d | PFC,  CeA,  NAc | Microarray |
| Smith et al., 2020 | 24 (PFC)  19 (CeA)  24 (NAc) | C57BL/6 Mice | CIE Vapor  (non-drinker) | 6 wk | 7 d | PFC,  CeA,  NAc | Microarray |
| Farris et al.,  2020 | 21 (PFC) | C57BL/6J Mice | CIE Vapor + limited access drinking | 5-8 wks | 1 wk | PFC | RNA-Seq |
| Farris et al.,  2020 | 17 (PFC) | C57BL/6J Mice | CIE Vapor + limited access drinking | 5-8 wks | 72 h | PFC | RNA-Seq |

**Suppl. Tab. 2**: Detailed information about included datasets and respective studies for meta-analysis in humans. PMI= post-mortem interval, BA= Brodmann Area, m= male, f= female. Included samples had a RNA integrity number (RIN) of >5. All studies included brain tissue from the NSWTRC at the University of Sydney, Australia, except Hade et al. (2021) who derived their tissue from the Estonian Forensic Science Institute, Tallinn, Estonia.

| *Study* | *n (AUD / Ctr)* | *Diagnosis* | *PMI (±SD) (hrs)* | *Brain Region* | *Method* |
| --- | --- | --- | --- | --- | --- |
| Liu et al., 2006 | 13m, 1f/ 10m, 3f | ≥80g alcohol per day | 28(±14) | Superior FC | Microarray |
| Ponomarev et al., 2012 | 17/ 15 | DSM-IV | not reported | Superior FC, BLA | Microarray |
| Manzardo et al., 2013 | 6m, 3f/ 6m, 3f | DSM-IV | 26.3 (±9.6) | PFC | Microarray |
| Wang et al., 2013 | 16m, 7f/ 16m, 7f | DSM-IV | not reported | BA9 | Microarray |
| Farris et al., 2015 | 16/ 16 | ≥7 yrs drinking,  DSM-IV | ≤48 | BA8 | RNA-Seq |
| Kapoor et al., 2019 | 51m, 14f/ 60m, 13f | DSM-IV | 33.66(±15.59)/  26.63(±13.25) | BA8 | RNA-Seq |
| Drake et al., 2020 | 34m, 7f/34m, 7f | - | 31.3(±12.1)/ 27.4(±12.5) | NAc | RNA-Seq |
| Rao et al., 2021 | 30/ 30 | DSM-IV | ≤48 | Superior FC, BLA, NAc | RNA-Seq |
| Hade et al., 2021 | 14/13 | alcohol-use-disorder-related diagnosis | 31.92(±12.88) | PFC | Microarray |
| Zillich et al., 2022 | 48/51 | DSM-IV | 37.07(±15.79)/  30.7 (±15.57) | NAc | RNA-Seq |

**Suppl. Tab. 3**: Detailed information about included datasets and respective studies for meta-analysis in monkeys.

| *Study* | *n* | *Strain* | *Treatment* | *Exposure length* | *Abstinence length* | *Brain Region* | *Method* |
| --- | --- | --- | --- | --- | --- | --- | --- |
| Bogenpohl et al., 2019 | 43 | Macaca mulatta | Drinking ad libitum | 1 yr | Up to 4 h | mPFC (BA24, 15, 32) | Microarray |
| Walter et al., 2020 | 23 | Macaca fascicularis | Drinking ad libitum | 6 m | None | PFC (BA46) | RNA-Seq |
| Fei et al. (unpublished) | 28 | Macaca mulatta | Drinking ad libitum | 646 d (avg.) | 4 wk | PFC (BA46) | RNA-Seq |
| Fei et al. (unpublished) | 24 | Macaca mulatta | Drinking ad libitum | 226 d (avg.) | None | PFC (BA46) | RNA-Seq |

**Suppl. Tab. 4** : Search terms and numbers of results for different rodent models of alcohol dependence. Studies described as suitable for quantitative analysis provide transcriptome wide data from RNA-Seq or Microarray Studies.

| **model** | **search terms** | **records identified through database search** | **titles screened** | **abstracts screened** | **full-text articles assessed** | **studies suitable for quantitative analysis** |
| --- | --- | --- | --- | --- | --- | --- |
| Reinstatement model | (alcohol OR ethanol) AND (rats OR mice OR rat OR mouse) AND (ribonucleic acid OR RNA OR RNA-seq OR mRNA OR ncRNA OR miRNA OR siRNA OR piRNA OR snRNA OR microRNA OR expression OR gene OR cDNA OR transcription OR transcript OR transcripts OR transcriptional OR transcriptomic OR translation OR transcriptome OR transcriptomics OR translation) AND ("alcohol-induced reinstatement" OR "cue-induced reinstatement" OR "cues-induced reinstatement" OR "priming" OR "stress-induced reinstatement" OR yohimbine OR "yohimbine-induced reinstatement" OR "foot-shock" OR "ABA" OR "renewal" OR "ABA renewal model" OR cue OR cues OR incubation OR extinction OR reinstatement) | 3124 | 3124 | 209 | 85 | 0 |
| ADE (alcohol deprivation effect) | (alcohol OR ethanol) AND (rats OR mice OR rat OR mouse) AND (ribonucleic acid OR RNA OR RNA-seq OR mRNA OR ncRNA OR miRNA OR siRNA OR piRNA OR snRNA OR microRNA OR expression OR gene OR cDNA OR transcription OR transcript OR transcripts OR transcriptional OR transcriptomic OR translation OR transcriptome OR transcriptomics OR translation) AND (ADE OR "alcohol deprivation effect" OR "alcohol-deprivation effect" OR "alcohol deprivation" OR "alcohol-deprivation" OR "ADEs" OR "ethanol deprivation" OR "ethanol-deprivation" OR "ethanol-deprivation effect" OR "ethanol-deprivation effect") | 76 | 76 | 69 | 68 | 1 |
| Postdependent | (alcohol OR ethanol) AND (rats OR mice OR rat OR mouse) AND (ribonucleic acid OR RNA OR RNA-seq OR mRNA OR ncRNA OR miRNA OR siRNA OR piRNA OR snRNA OR microRNA OR expression OR gene OR cDNA OR transcription OR transcript OR transcripts OR transcriptional OR transcriptomic OR translation OR transcriptome OR transcriptomics OR translation) AND (post-dependent OR "post dependent" OR postdependent OR cie OR "chronic intermittent"). For human post-mortem studies, we used the following keywords: (“post-mortem” OR “postmortem” OR “human” OR “patients”) AND (“brain”) AND (“alcohol” OR “alcoholic” OR “alcohol dependence” OR “alcoholism” OR “AUD” OR “alcohol use disorder” OR “alcohol addiction”) AND (“transcriptome” OR “transcriptomic” OR “transcriptomics” OR “gene-expression” OR "gene expression" OR “mRNA” OR “messenger RNA”) | 656 | 534 | 227 | 179 | 5 (= 8 datasets) |
| Alcohol preferring lines | (alcohol OR ethanol) AND ("alcohol preferring" OR "alcohol-preferring" OR "ethanol preferring" OR "ethanol-preferring" OR "genetically selected" OR "msP rats" OR "AA rats" OR "ANA rats" OR "HAD rats" OR "LAD rats" OR "Indiana P rats" OR "P rats" OR "sP rats" OR "sNP rats" OR "UChB rats" OR "UChA rats" OR "iP rats") AND (ribonucleic acid OR RNA OR RNA-seq OR mRNA OR ncRNA OR miRNA OR siRNA OR piRNA OR snRNA OR microRNA OR expression OR gene OR cDNA OR transcription OR transcript OR transcripts OR transcriptional OR transcriptomic OR translation OR transcriptome OR transcriptomics OR translation) NOT (systematic review[pt] OR meta-analysis[pt] OR review[pt]) | 308 | 308 | 285 | 217 | 17; several diff. lines; brain regions very variable |
| Aversion resistent | (alcohol OR ethanol) AND (rats OR mice OR rat OR mouse) AND („aversion-resistant“ OR „aversion resistant“ OR „aversion-resistant drinking“ OR „aversion resistant drinking“ OR „aversion-resistant intake“ OR „aversion resistant intake“ OR „aversion-resistant consumption“ OR „aversion resistant alcohol consumption“) | 43 | 43 | 29 | 28 | 0 |

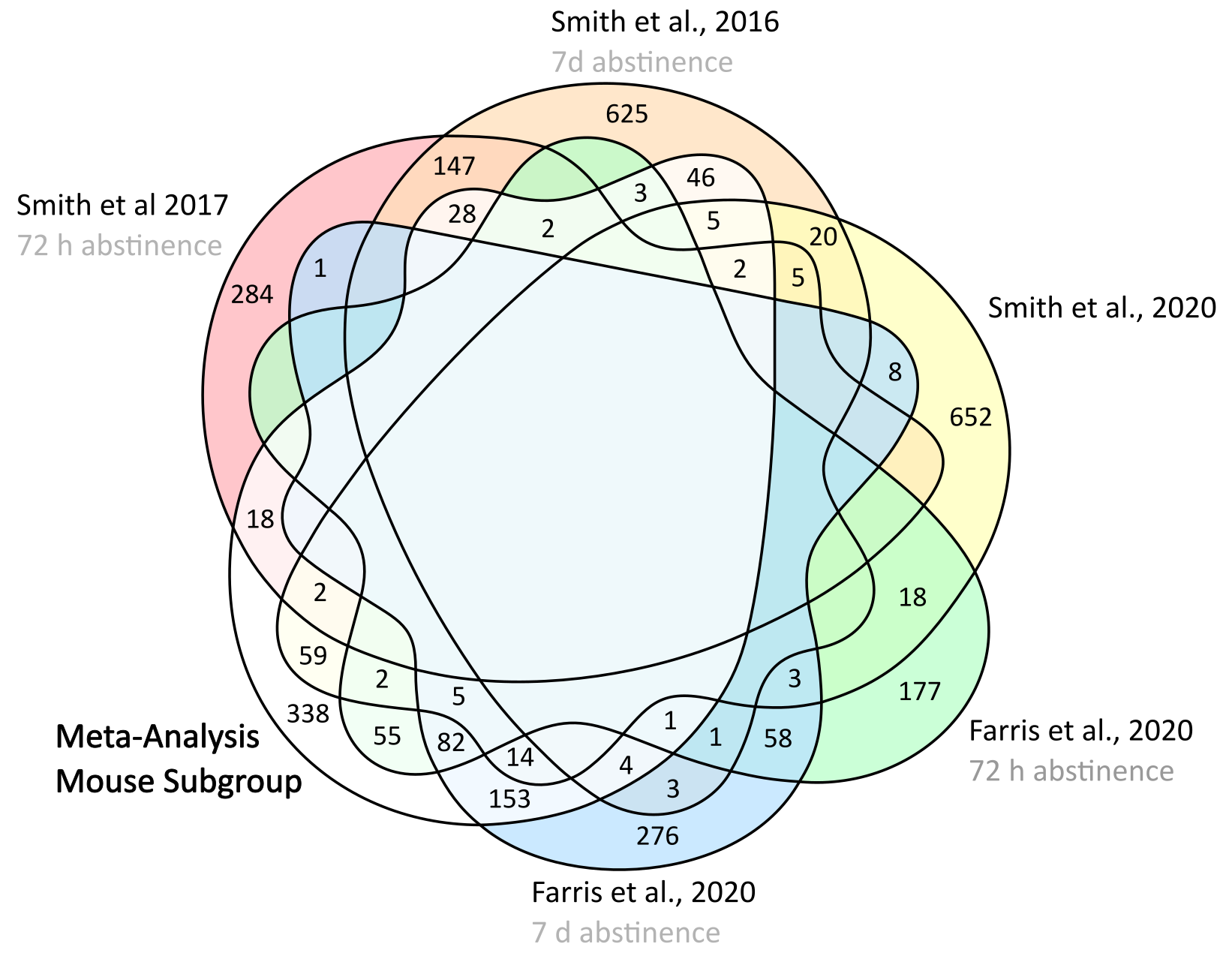

**Suppl. Fig. 1:** Venn Diagram comparing overlap of mouse **PFC** meta-analysis with original studies focusing on mouse brain tissue.

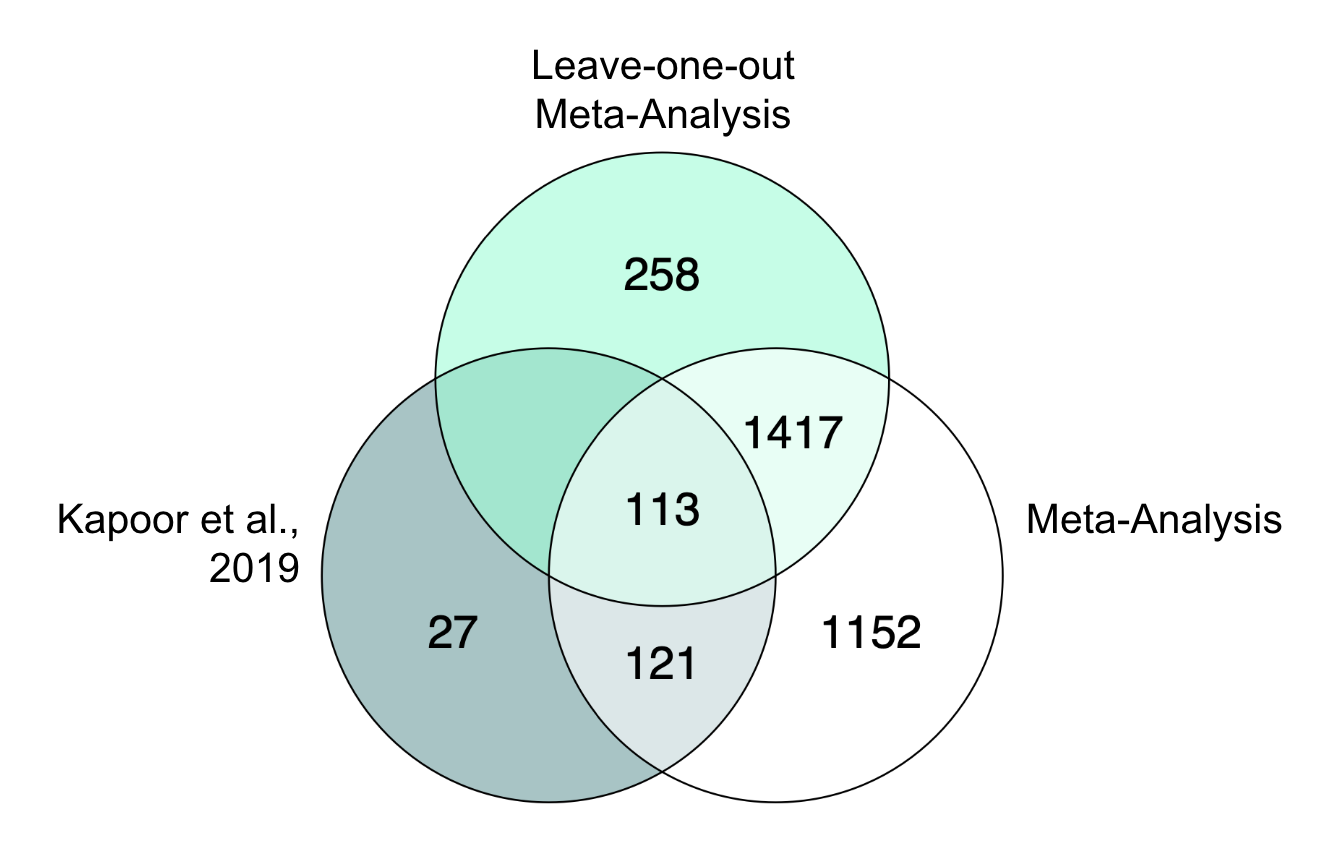

**Suppl. Fig. 2**: Venn Diagram representing leave-one-out meta-analysis in comparison to whole human PFC meta-analysis. LOO analysis was performed to provide further insights to the potential bias that could have been introduced by the highly varying number of individuals per original study. Therefore, the study with the highest number of individuals was excluded in LOO and the results were compared to the human meta-analysis. Since the results from LOO are mostly a subset of the final meta-analysis results, we assume that Kapoor et al. increased the detection capability of the meta-analysis without introducing substantial bias.

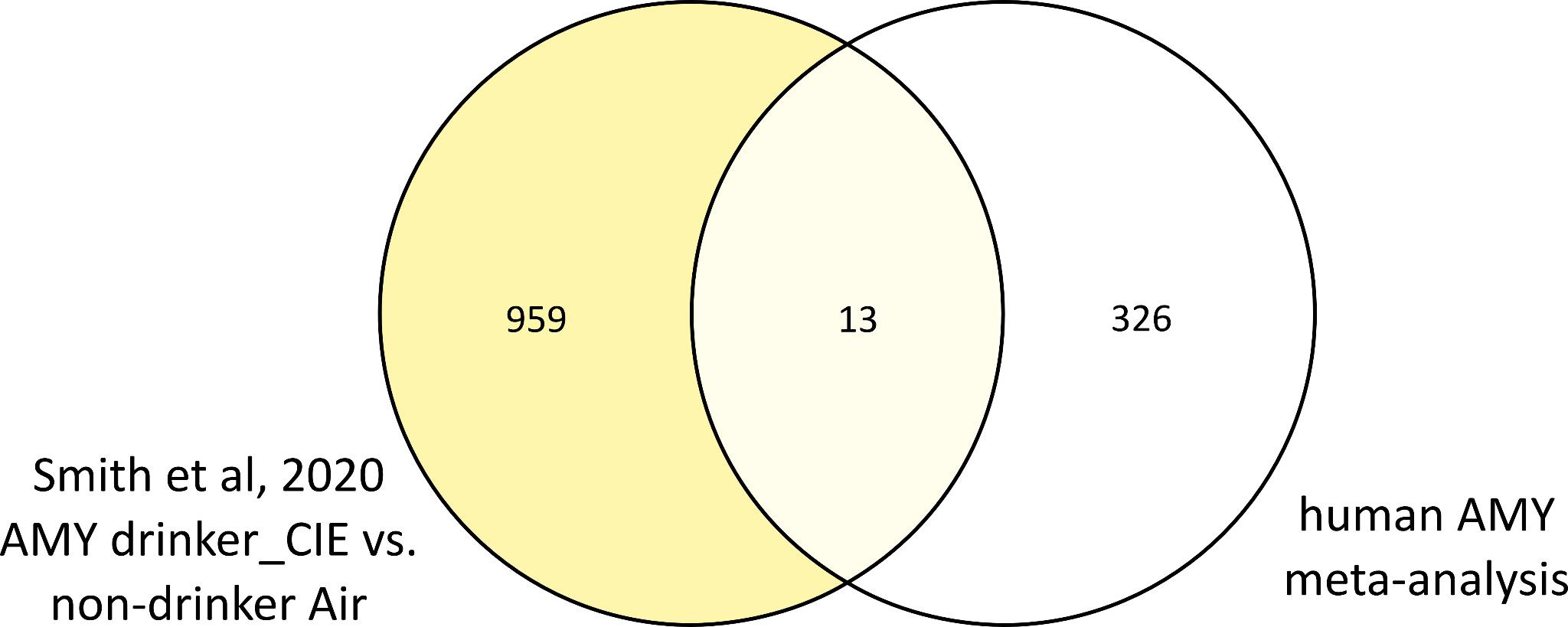

**Suppl. Fig 3: Overlapping results of human AMY Meta-analysis and Smith et al., 2020 Drinker CIE vs non-drinker Air group** Meta-analysis of human postmortem tissue from amygdala resulted in 339 DEGs (FDR < 0.05). Analysis of drinker CIE mice vs. non-drinker air-exposed mice by Smith et al., 2020 yielded 972 DEGs (FDR < 0.05). As the latter might be a better model of the human scenario than only CIE exposed mice, results were compared to the results of human meta-analysis. 13 genes overlap, namely "AXIN1"	"BCL7C"	"DRAP1"	"GFPT2"	"IFITM1" "MFHAS1" "MYO9B"	"PREB"	"SAFB2"	"SDC4"	"SF3B2"	"SLC44A2" "TRPS1".

**References**

Bogenpohl, J. W., Smith, M. L., Farris, S. P., Dumur, C. I., Lopez, M. F., Becker, H. C., Grant, K. A., & Miles, M. F. (2019). Cross-species co-analysis of prefrontal cortex chronic ethanol transcriptome responses in mice and monkeys. *Frontiers in Molecular Neuroscience*, 197.

Drake, J., McMichael, G. O., Vornholt, E. S., Cresswell, K., Williamson, V., Chatzinakos, C., Mamdani, M., Hariharan, S., Kendler, K. S., Kalsi, G., Riley, B. P., Dozmorov, M., Miles, M. F., Bacanu, S.-A., & Vladimirov, V. I. (2020). Assessing the Role of Long Noncoding RNA in Nucleus Accumbens in Subjects With Alcohol Dependence. *Alcoholism: Clinical and Experimental Research*, *44*(12), 2468-2480. https://doi.org/https://doi.org/10.1111/acer.14479

Farris, S. P., Arasappan, D., Hunicke-Smith, S., Harris, R. A., & Mayfield, R. D. (2015). Transcriptome organization for chronic alcohol abuse in human brain. *Molecular psychiatry*, *20*(11), 1438-1447.

Farris, S. P., Tiwari, G. R., Ponomareva, O., Lopez, M. F., Mayfield, R. D., & Becker, H. C. (2020). Transcriptome analysis of alcohol drinking in non-dependent and dependent mice following repeated cycles of forced swim stress exposure. *Brain sciences*, *10*(5), 275.

Hade, A.-C., Philips, M.-A., Reimann, E., Jagomäe, T., Eskla, K.-L., Traks, T., Prans, E., Kõks, S., Vasar, E., & Väli, M. (2021). Chronic alcohol use induces molecular genetic changes in the dorsomedial thalamus of people with alcohol-related disorders. *Brain sciences*, *11*(4), 435.

Kapoor, M., Wang, J. C., Farris, S. P., Liu, Y., McClintick, J., Gupta, I., Meyers, J. L., Bertelsen, S., Chao, M., Nurnberger, J., Tischfield, J., Harari, O., Zeran, L., Hesselbrock, V., Bauer, L., Raj, T., Porjesz, B., Agrawal, A., Foroud, T., . . . Goate, A. (2019). Analysis of whole genome-transcriptomic organization in brain to identify genes associated with alcoholism. *Transl Psychiatry*, *9*(1), 89. https://doi.org/10.1038/s41398-019-0384-y

Liu, J., Lewohl, J. M., Harris, R. A., Iyer, V. R., Dodd, P. R., Randall, P. K., & Mayfield, R. D. (2006). Patterns of gene expression in the frontal cortex discriminate alcoholic from nonalcoholic individuals. *Neuropsychopharmacology*, *31*(7), 1574-1582. https://doi.org/10.1038/sj.npp.1300947

Manzardo, A. M., Gunewardena, S., Wang, K., & Butler, M. G. (2014). Exon microarray analysis of human dorsolateral prefrontal cortex in alcoholism. *Alcohol Clin Exp Res*, *38*(6), 1594-1601. https://doi.org/10.1111/acer.12429

Meinhardt, M. W., Hansson, A. C., Perreau-Lenz, S., Bauder-Wenz, C., Stählin, O., Heilig, M., Harper, C., Drescher, K. U., Spanagel, R., & Sommer, W. H. (2013). Rescue of infralimbic mGluR2 deficit restores control over drug-seeking behavior in alcohol dependence. *Journal of Neuroscience*, *33*(7), 2794-2806.

Osterndorff-Kahanek, E. A., Becker, H. C., Lopez, M. F., Farris, S. P., Tiwari, G. R., Nunez, Y. O., Harris, R. A., & Mayfield, R. D. (2015). Chronic ethanol exposure produces time-and brain region-dependent changes in gene coexpression networks. *PLoS One*, *10*(3), e0121522.

Ponomarev, I., Wang, S., Zhang, L., Harris, R. A., & Mayfield, R. D. (2012). Gene coexpression networks in human brain identify epigenetic modifications in alcohol dependence. *J Neurosci*, *32*(5), 1884-1897. https://doi.org/10.1523/jneurosci.3136-11.2012

Rao, X., Thapa, K. S., Chen, A. B., Lin, H., Gao, H., Reiter, J. L., Hargreaves, K. A., Ipe, J., Lai, D., Xuei, X., Wang, Y., Gu, H., Kapoor, M., Farris, S. P., Tischfield, J., Foroud, T., Goate, A. M., Skaar, T. C., Mayfield, R. D., . . . Liu, Y. (2021). Allele-specific expression and high-throughput reporter assay reveal functional genetic variants associated with alcohol use disorders. *Mol Psychiatry*, *26*(4), 1142-1151. https://doi.org/10.1038/s41380-019-0508-z

Smith, M. L., Lopez, M. F., Archer, K. J., Wolen, A. R., Becker, H. C., & Miles, M. F. (2016). Time-course analysis of brain regional expression network responses to chronic intermittent ethanol and withdrawal: implications for mechanisms underlying excessive ethanol consumption. *PLoS One*, *11*(1), e0146257.

Smith, M. L., Lopez, M. F., Wolen, A. R., Becker, H. C., & Miles, M. F. (2020). Brain regional gene expression network analysis identifies unique interactions between chronic ethanol exposure and consumption. *PLoS One*, *15*(5), e0233319.

Walter, N. A., Zheng, C. L., Searles, R. P., McWeeney, S. K., Grant, K. A., & Hitzemann, R. (2020). Chronic voluntary ethanol drinking in cynomolgus macaques elicits gene expression changes in prefrontal cortical area 46. *Alcoholism: Clinical and Experimental Research*, *44*(2), 470-478.

Wang, F., Gelernter, J., & Zhang, H. (2013). Differential Expression of miR-130a in Postmortem Prefrontal Cortex of Subjects with Alcohol Use Disorders. *J Addict Res Ther*, *4*(155). https://doi.org/10.4172/2155-6105.1000155

Zillich, L., Poisel, E., Frank, J., Foo, J. C., Friske, M. M., Streit, F., Sirignano, L., Heilmann-Heimbach, S., Heimbach, A., Hoffmann, P., Degenhardt, F., Hansson, A. C., Bakalkin, G., Nöthen, M. M., Rietschel, M., Spanagel, R., & Witt, S. H. (2022). Multi-omics signatures of alcohol use disorder in the dorsal and ventral striatum. *Translational Psychiatry*, *12*(1), 190. https://doi.org/10.1038/s41398-022-01959-1
